# Construction and Immunogenicity Detection of Canine Parvovirus-like Particles Fusion with Canine Febrile Antigen Epitopes

**DOI:** 10.64898/2026.01.18.700204

**Authors:** Yuhan Zhu, Xintao Liu, Ying Xu, Guozhen Zhang, Yuhe Yin, Congmei Wu

## Abstract

This study aimed to design and evaluate the immunogenicity of a dual-valent virus-like particles (VLP) vaccine that can simultaneously target Canine distemper virus (CDV) and Canine parvovirus virus (CPV).By bioinformatic analysis, conserved antigen epitopes of the three major functional proteins of CDV were screened and inserted into CPV-VP2 proteins by two different methods to construct recombinant expression plasmids CDPV1 versus CDPV2.Three-dimensional structure, hydrophobicity and stability predictions of the two recombinant proteins showed that their hydrophobicity were 0.233 and 0.251, and their structural stability scores were 0.77 and 0.78, respectively.Recombinant plasmids were co-transformed with molecular chaperone pTf16 to be expressed in E. coli BL21(DE3), respectively, and the optimal expression conditions were determined after optimization: 0.25 mmol/L IPTG, 2 g/L L-arabinose in duction, and culture at 25°C for 16 h.Purified recombinant proteins can self-assemble in vitro to form VLPs about 23.5 nm in diameter with a hemagglutination titer of 1:2^9^.The mouse immune test showed that the hemagglutination inhibition titer peaked on the 7th day after the third immunization, and the neutralizing Antibody level could reach up to 1:2^8^.The CPV VLP s constructed in this institute carrying CDV antigen epitopes were able to successfully assemble in vitro and were well immunogenic, providing experimental basis for the development of a canine polyvalent vaccine.

## Introduction

Canine distemper virus ( CDV ) is the main canine pathogen of the genus measles virus in the family Canine distemper viruses, it is a highly contagious, highly lethal multi-system viral disease. The natural hosts of infection are both canine and anthropozoon animals^[1].^Symptoms such as vomiting, pneumonia, diarrhea, dehydration, and anorexia occur in the early stages of infection[2].Among them, N protein (nucleocapsid protein), F protein (fusion protein), and H protein (haemagglutinin protein) are the main structural proteins of CDV virus, which p lay a key role in the virus’s infection mechanism, host interaction, and immune response ^[3]^.Neutralizing epitopes of CDVs are found predominantly on N and F proteins, which can elicit a strong immune response and neutralize Antibodies ^[4]^.While the H protein of CDV, as a glycoprotein of the viral surface recognition receptor, affects the antigenicity and host recognition of CDV, the domain of H protein has been shown to play a key role in the cross-species transmission of CDV ^[5]^.

Canine parvovirus virus ( CPV ) is the main viral cause of acute canine enteritis, clinical symptoms including fever, leukopenia, diarrhea, dehydration and anorexia.CPVs were highly contagious and mortality rates were also high .Among these, VP1 protein (secondary capsid protein), VP2 protein (major capsid protein), and VP3 protein (capsid-associated protein) are the major structural proteins of the virus, and CPV-VP2, the most abundant protein found on the CPV capsid, is the major structural unit that constitutes the viral capsid, and its N-terminal domain has been shown to be an excellent target for the development of synthetic peptide vaccines. VP2 consists of 584 amino acid residues, forming eight antiparallel β-folded and 4-loop structures that surround each other to form a triple fibrin that contains the major epitope of VP2 protein and the binding site of the virus to the host receptor ^[6]^.In recent years, VP2 protein has gained attention as a vaccine for virus-like granule protein and there are many related vaccines used in animals and humans.

Virus-like particles (VLPs) retain the antigen epitope and structural morphology of the natural viral particles and effectively present glycoproteins and other antigen components on the surface of the viral particles ^[7]^, while VLPs can induce a strong cellular immune response ^[ 8]^.VLPs allow for small replacement or insertion of exogenous antigens and do not affect the assembly of VLPs, thus, VLPs can be used as Metastasis vectors carrying exogenous protein or antigen epitopes to produce chimeric VLPs for bivalent or polyvalent vaccines ^[9]^.

This study aims to combine the antigen epitope gene of CDV with the CPV-VP2 gene to form chimeric CPV-VLPs, which enable these antigens to possess both properties, and to explore the feasibility of chimeric VLPs using E. coli system expression.

## Materials and methods

### Ethics Statement

Experimental studies were approved by the Animal Ethics Committee of Changchun Longsheng Experimental Animal Technology Co., Ltd. (Ethics Review No: CCLSLL-2025082817).The license for use in experimental animals is: SYXK (Gi) 2022-008.The license for the pro duction of experimental animals is: SCXK (Beijing) 2024-0001.), experimental operations were followed the China Guideline on the Welfare and Ethics of Laboratory Animals.

### Experimental Animals

Thirty healthy female BALB/c mice, 4 weeks of age, weighing 18-20 g, were purchased from Beijing Sbei Fu Biotechnology Co., Ltd. The breeding conditions are as follows: Chang chun Xinuo Co., Ltd. Clean level animal room (barrier system), room temperature 25 °C, relative humidity 55%, alternating light 12 h bright and 12 h dark, and using independent ventilation cages (IVC) for breeding. All cages and pads were used after autoclaving, and the mice freely ingested drinking water. The experimental animals and procedures involved in the study were approved by the Ethics Committee and strictly adhered to the rules and regulations of animal experiments. Before the beginning of the experiment, all investigators received training on animal care and procedures. During the experiment, to minimize animal pain and injury, we paid attention to the diet and activity of the mice in a timely manner. Each mouse under went tail venous plexus blood sampling every 7 days, 150 μL of peripheral blood sample was collected, the blood sample was coagulated in a centrifuge tube for 1 h, centrifuged at 400 0 rpm for 10 min, Serum was collected and heat-inactivated at 56 °C for 30 min, and then stored at -80 °C for analysis. At 42d, mice were euthanized by anesthesia injection (1% pentobarbital sodium solution, 45 mg/kg). All procedures were in accordance with animal ethics-related norms.

### Plasmids, Species, and Molecular Companions

Recombinant plasmids CDPV1, CDPV2 were synthesized by Nanjing Jinsrui Biotechnology Co., Ltd.; E.coli BL21 (DE3) competent cells were purchased from Beijing All-in-One Gold Biotechnology Co., Ltd.; molecular chaperones pGro7, pTf16 were purchased from TaKaRa Co., Ltd.

### Main reagent

The primary antibody (His Tag Mouse Monoclonal Antibody), the secondary antibody ( HRP-labelled goat anti-mouse IgG), and the DAB horseradish peroxidase chromogenic kit use d in Western Blot were purchased from Shanghai Biyuntian Biotechnology Co., Ltd.; isopropyl-β-d-thiogalactosidase (IPTG), and L-arabinose were purchased from Shanghai McClin Biochemical Technology Co., Ltd.; and the His tag protein affinity chromatography pre-assembled column (Ni2 column) was purchased from Detai Biotechnology Co., Ltd.

### Construction and Expression of Recombinant Plasmids C DPV1 and CDPV2

To successfully express recombinant proteins in E.coli, this trial predicted and screened the antigen epitopes of CDV and connected the screened antigen epitopes to CPV VP2. The resulting complete sequence was synthesized by Nanjing Jinsrui Biotechnology Co., Ltd.Using E. coli BL21 (DE3) competent cells, CDPV1 and CDPV2 plasmids were converted into CDPV2 plasmids to obtain CDPV1 and CDPV2 bacteria. The bacteria were evenly coated with LB solid media containing kanamycin (Kan) resistance for culture. After culture, the monoclonal strains were selected and seeded in 50 mL LB liquid medium containing Kan resistance f or oscillating culture at 37 °C at 220 r/min; when the OD600 value reached 0.6-0.8, IPTG at a final concentration of 0. 25 mM/L was added and induced at 25 °C for 16 h; the induced bacteria were collected for US fragmentation, and then SDS-PAGE and Western blot were performed to analyze the expression products.

### Construction of two molecular chaperones coexpressings trains and soluble expression of recombinant proteins

The molecular chaperones pGro7, pTf16 were co-transformed with CDPV1 and CDPV2 and treated in competent cells. The co-expressed strains were co-cultured on LB solid media with dual antibodies to kanamycin (Kan) and chloramphenicol (Cm); incubated overnight at 3 7°C, and then single colonies were picked and seeded in LB liquid medium at 37°C, 220 r/m in, and incubated until the absorbance OD600 value reached 0.6-0.8 with L-arabinose and IPTG.After induction of expression at 25 °C, 220 r/min for 14 h, the resulting induction bacteria US was fragmented, identified by SDS-PAGE and the expression products were analyzed.

### Optimization of expression conditions of recombinant proteins

Exploring the induction temperature of the inducible protein and the concentration of IPTG added at the time of induction.18, 25, 30 °C was selected as the induction temperature, and expression was induced with 0. 25 mmol/L IPTG, 2 g/L L-arabinose for 14 h. After US crushing, SDS-PAGE was performed for identification, and the optimal induction temperature was selected.0.25, 0.5mmol/L were selected as IPTG concentrations, which were induced to express for 14 h at the optimal induction temperature, and identified by SDS-PAGE after US fragmentation to select the optimal induction IPTG concentration.

### Purification of recombinant expressed proteins

The bacteria obtained by optimal induction conditions were US broken up, and the collected US supernatants were filtered through 0.45μm filters, purified by affinity chromatograph y using a Ni2 column. After equilibration, loading, washing, and elution (using imidazol elution gradient), the purified protein was obtained. The purity of the purified protein was detected by SDS-PAGE and Western blot.

### Self-assembly and Identification of CDPV1 and CDPV2

Purified CDPV1 and CDPV2 were placed in dialysis bags, treated with Tris buffer (pH= 8.0) and NaCl assembly solution (at a concentration of 250 mmol/L) at 4°C, and the buffer and assembly solution were changed every 4 hours for 4 repeats to complete self-assembly.Samples of self-assembled CDPV1 versus CDPV2 were measured for morphology and particle size using transmission electron microscopy (TEM) and dynamic light scatter (DLS), respectively.

### Determination of the Coagulation Values of CDPV1 and CDPV2

Coagulation tests were performed on CDPV1, CDPV2 to determine the hemagglutination potency of VLPs. The 96-well plate was filled with 25 μL of CDPV1 and CDPV2, which was fully mixed and diluted to the last well by a 2-fold ratio. Finally, 25 μL was aspirated and discarded. 50 μL of 1% porcine red blood cell, RBC suspension was added to each well. The results were determined after being left at room temperature for 1 h. Finally, the highest dilution of 1% porcine red blood cell, RBC suspension could be used as the coagulation titer of the sample.

### Animal immunization protocol

Thirty 4-week-old BALB/c female mice were randomly divided into six groups, each with a first immunization on day 0, booster immunization on days 7, and 14, Serum was collected every 7 days, heat-inactivated for 30 min at 56°C, and then stored at -80 °C for analysis, and mice were sacrificed on day 42 as shown in Figure 1.The immunization was performed by intramuscular injection with a protein immunization group: CDV, CPV double-inactivated vaccine group, CDPV1 group, CDPV2 group, CDPV1 Al(OH)3 adjuvant, CDPV2 Al(OH)3 adjuvant group, and PBS control arm. The recombinant protein dose of the immunized group was 20 μg per mouse, in which Al(OH)_3_ adjuvants were mixed with recombinant protein in a 1:1 ratio to prepare the vaccine.PBS was diluted to 100 μL as shown in Table 1.All animals died as a result of euthanasia after the end of the trial.

**Table 1.** Experimental grouping and immunization dosage.

| Immunogen | Injection volume (μL/mouse) | Antigen Dose (μg) | Immune pathway | Immune times |
| --- | --- | --- | --- | --- |
| PBS | 100 | / | Injection of muscle | 3 |
| double inactivated vaccine | 100 | / | Injection of muscle | 3 |
| CDPV1 | 100 | 20 | Injection of muscle | 3 |
| CDPV2 | 100 | 20 | Injection of muscle | 3 |
| CDPV1+Al(OH)<sub>3</sub> | 100 | 20 | Injection of muscle | 3 |
| CDPV2+Al(OH)<sub>3</sub> | 100 | 20 | Injection of muscle | 3 |

**Figure 1.**
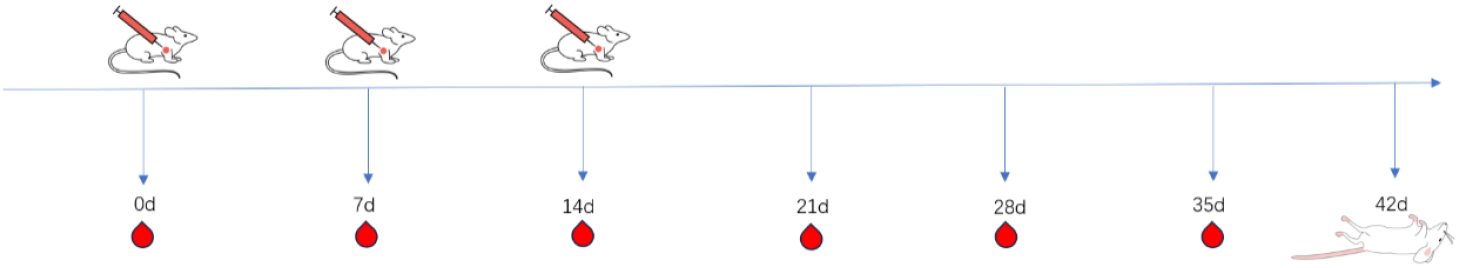
Schematic diagram of the immunization schedule.

### Determination of the potency of hemagglutination inhibition

96-well plate, 25μL PBS was added to the plate; 25μL of Serum for testing was added to the first well, and 2 wells were repeated; the sample in the first well was repeatedly puffed with PBS 5 times with a pipette, and fully mixed; 25μL of liquid was aspirated from the first well to the second well, and step 3 was repeated until the last well, and the 25μL of liquid aspirated from the last well was discarded; 25 μL of 8 units of hemagglutinin was added to all wells, which were allowed to operate at 37°C for 30 min; 50 μL of 1% porcine red blood cell, RBC suspension was added to all wells, which was gently mixed and placed in the refrigerator at 2 to 8 °C for 1 h; negative control holes were placed on the same plate; they were removed after 1 h of reaction at 2 to 8 °C, and the plate reads were determined by stan ding up. The highest dilution of complete agglutination is used as the result of the blood clot assay.

### Detection of Neutralizing Antibodies

The harvested mouse Serum was filtered and sterilized with a 0.22 μm filter and diluted 10 times with maintenance medium followed by serial fold-dip dilution in 96-well plates.Mixed with an equal volume of 100 tissue half-infected amounts (TCID50) of CDV and CPV and cultured in a 5% CO_2_ incubator at 37°C for 1 h. Subsequently, 100 μL of the mixture Metastasis was added to FK81 cells in 96-well plates for 1 h in a 5% CO_2_ incubator at 37°C, followed by discarding the liquid in the 96-well plates, washing three times with PBS, and adding DMEM with 2% FBS. Each Serum sample was analyzed three times. After 48-72h of incubation, visualization of the cytopathic effect (CPE) was used to calculate TCID50 values

## Results

### Construction of recombinant plasmid CDPV1 and CDPV 2

#### Sequence conservation prediction

Antigenic epitopes of F, N, and H proteins were selected for fusion expression with the VP2 gene.By reviewing the relevant literature, we selected the major antigen epitopes identified in previous studies for CDV, mainly one B cell epitope and one CTL epitope for CDV-N proteins, one Th cell epitope, one T cell epitope, and one B cell epitope for F proteins. The major antigen epitopes are shown in Table 2.Conservative prediction of antigen epitopes was performed using the website IEDB Analysis Resource (immuneepitope.org) and results are shown in Table 3.

**Table 2.**
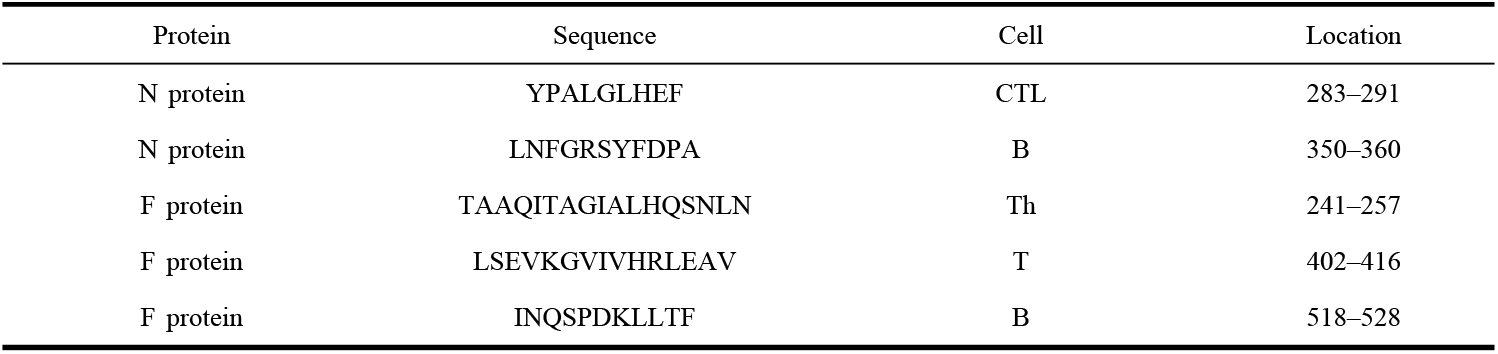
CDV major antigen epitopes.

**Table 3.**
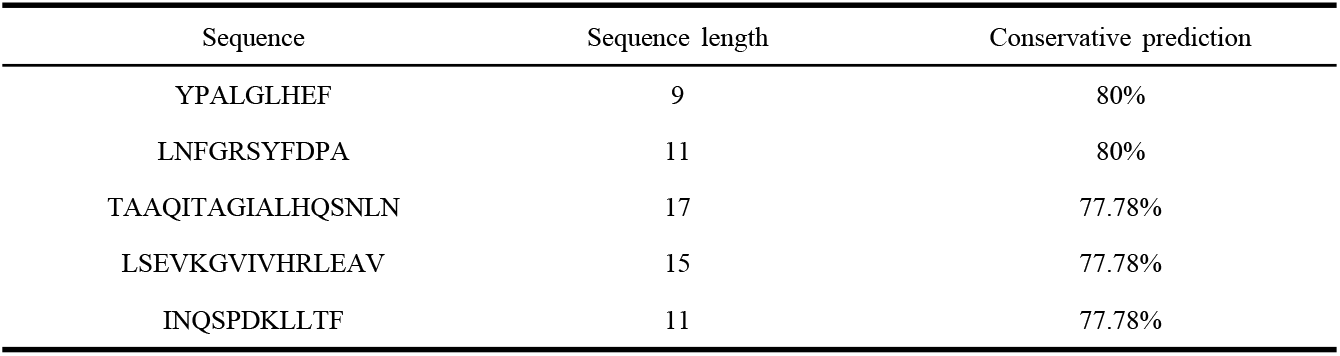
Conservative prediction of CDV major antigen epitopes.

#### Prediction of CDV H protein B cell epitopes

Prediction screening for antigen epitopes in the H protein region of canine distemper virus.Screening was performed by predicting through the IEDB Analysis Resource (immuneepitope.org) website, and the results are shown in Table 4. Ten epitopes were obtained, and further studies were performed through two-dimensional structure predictions. Because of the large number of beta folds and irregular coiling in the top-ranked sequence, the second-ranked sequence was selected for further study, and the results are shown in Figure 2.

**Table 4.**
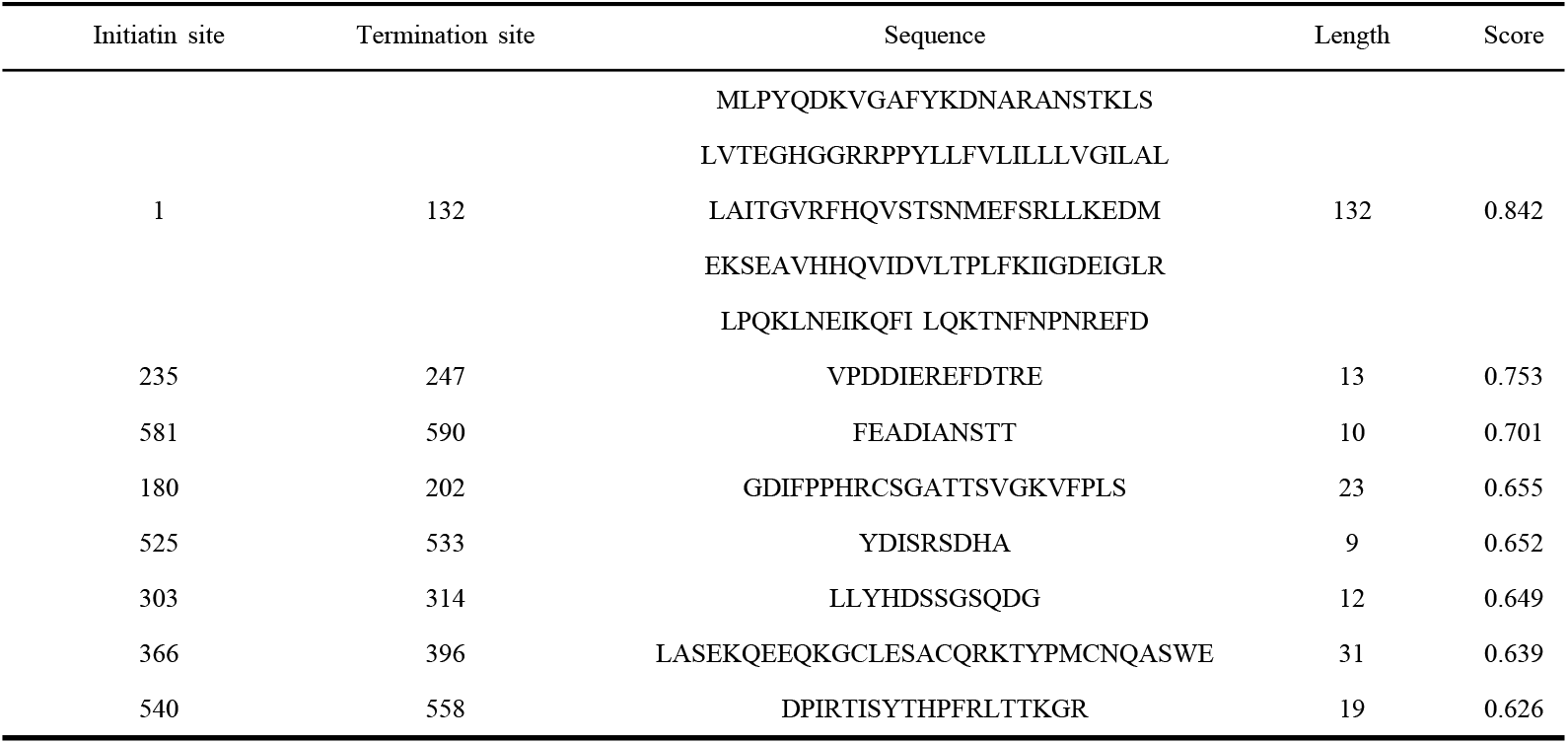
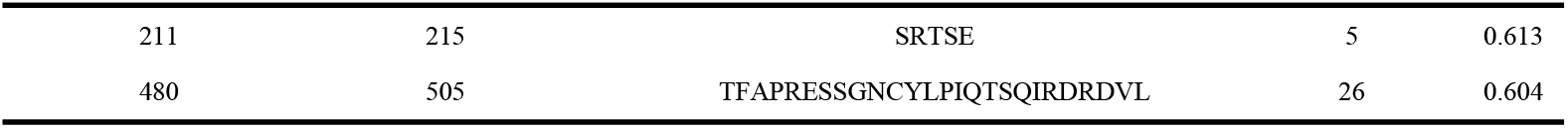
Main antigenic epitopes of CDV H protein.

**Figure 2.**
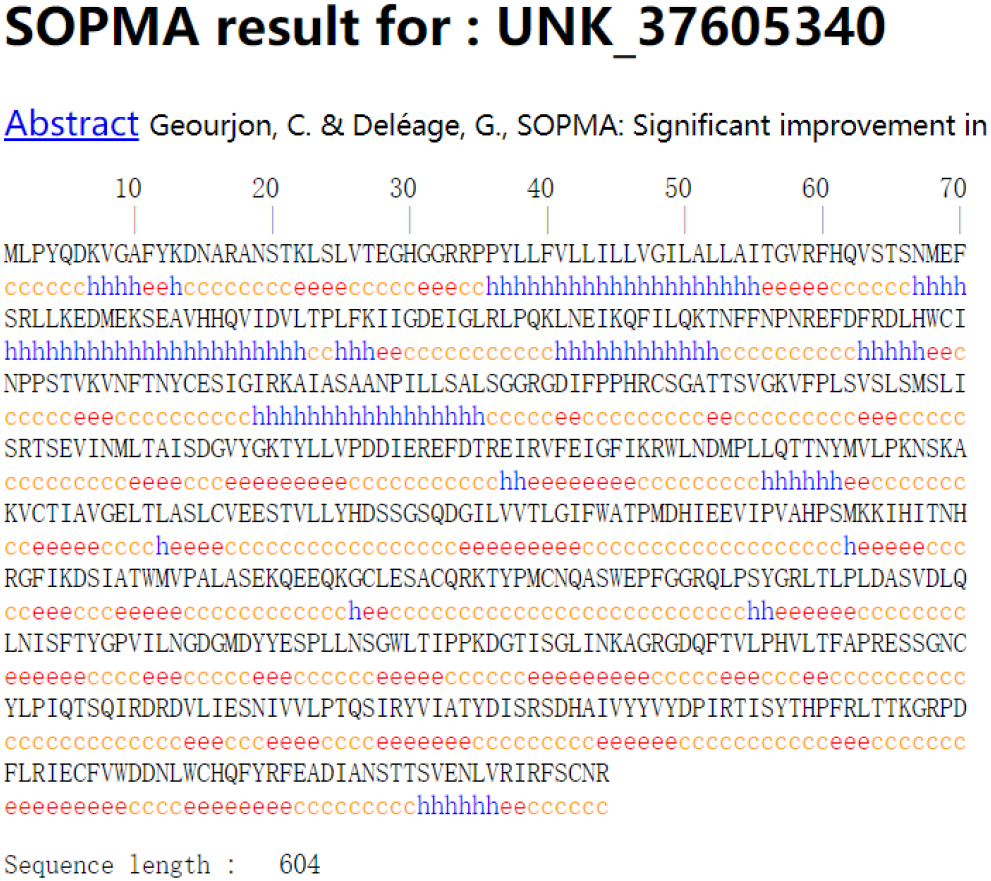
**Two-dimensional structure prediction of H protein**, h: α-helix; t: β-turn; c: irregular coiling; e: extended strand

#### Structure and prediction of CDPV1 and CDPV2

An antigenic epitope of CDV was inserted at the N-terminus of CPV-VP2, as shown in Fig. 3A, to name this recombinant plasmid CDPV1.The neutralizing epitope of CDV-H protein, the Th-T-B epitope of CDV-F protein, and the CTL-B epitope of CDV-N protein inserted at the N terminus of CPV-VP2, as shown in Fig. 3B, have been named CDPV2 for this recombinant protein.Predicting the hydrophobicity of the two expression proteins (ProtScale analysis (expasy.org) ) and the three-dimensional structure (Figs. 4-5) resulted in a target protein size of approximately 74kDa, soluble expression predicted by 0.233 (Fig. 4A), and three-dimensional structure predicted by 0.77 (Fig. 5A) and a target protein size of approximately 76kDa, soluble expression predicted by 0.251 (Fig. 4B), and three-dimensional structure predicted by 0.78 (Fig. 5B).The predicted results suggest that CDPV1,CDPV2 have a stable structure.

**Figure 3.**
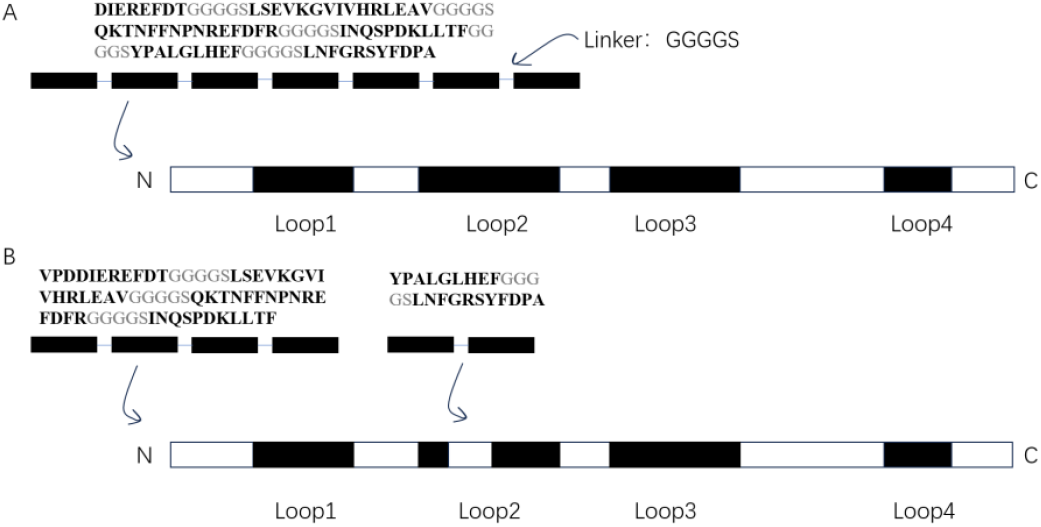
Schematic representation of the structure of the recombinant plasmid. A: CDPV1;B:CDPV2

**Figure 4.**
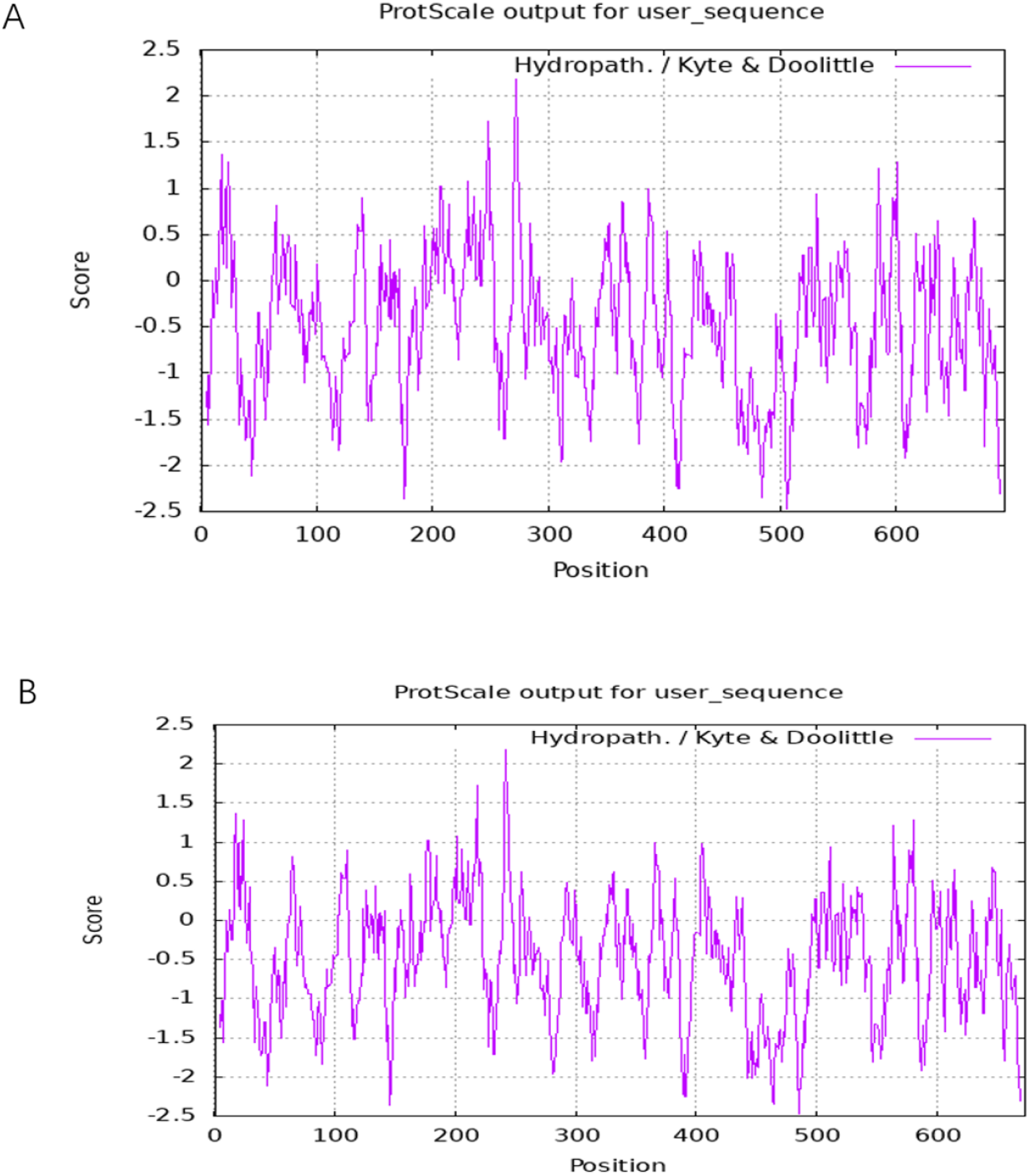
**Prediction results of hydrophobicity of CDPV1 and CDPV2**, A: CDPV1; B: CDPV2

**Figure 5.**
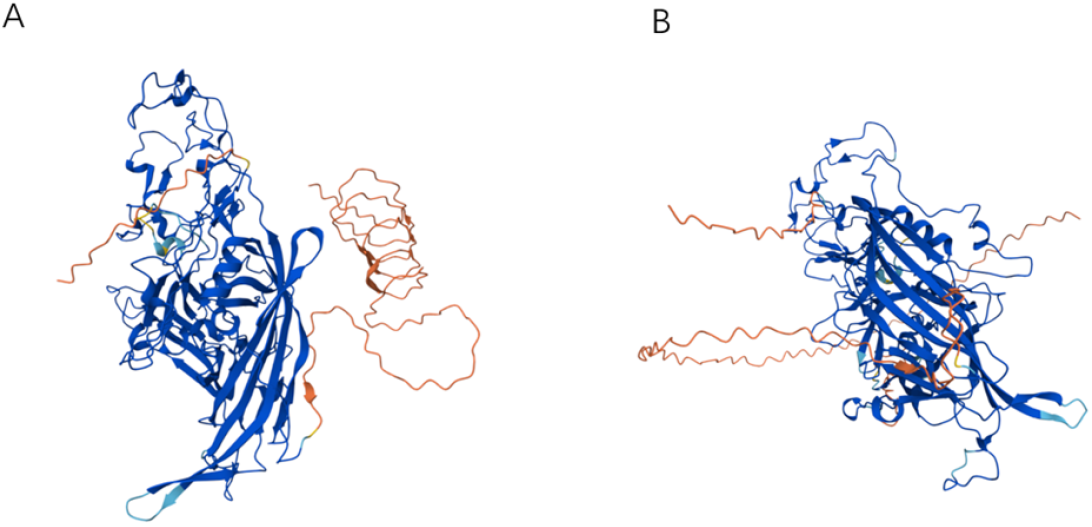
**Schematic diagram of three-dimensional structure of CDPV1 and CDPV2**, A: CDPV1; B: CDPV2

**Figure 6.**
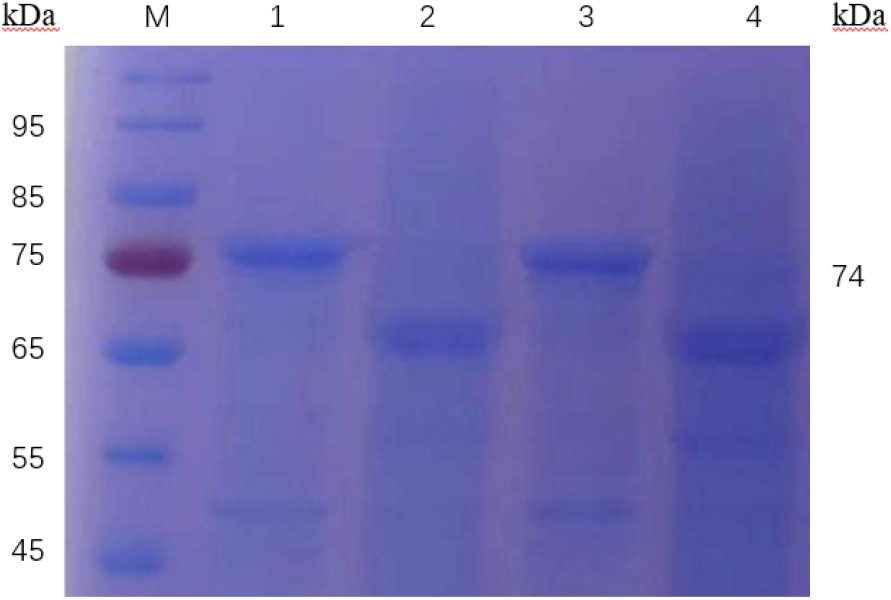
SDS-PAGE identification results of CDPV1, CDPV2. M: Maker; 1: CDPV2 precipitation; 2: CDPV2 supernatant; 3: CDPV1 precipitation; 4: CDPV1 supernatant

### Expression and purification of recombinant proteins

#### Expression of recombinant protein C DPV1 and CDPV2

Recombinant plasmids were converted separately into E.coli BL21(DE3) competent cells and induced to express by 0.5 mmol/L of IPTG at 25 °C.The SDS-PAGE analysis of the bacterial juice after US fragmentation showed that protein C DPV1 and CDPV2 expressed target proteins of expected size at 74kDa and 76kDa, respectively, but both were inclusion-some, s o we considered adding molecular chaperones for coexpression.

#### Recombinant Protein CDPV1, CDPV2 coexpressed with two molecular chaperones

CDPV1 and CDPV2 were converted to competent cells with two molecular chaperones, pGro7 and pTf16, respectively, and the recombinant bacteria were named CDPV1-pGro7, CDPV1-pTf16, CDPV2-pGro7, and CDPV2-pTf16.In the culture fluid of all 4 strains, L-arabinos eat a final concentration of 2mg/mL was added, and the expression conditions were screened and optimized.The results in Fig.7 show that CDPV1 coexpression with the molecular chaperone pGro7 at different temperatures is either inclusion-body expression and therefore is not selected for coexpression with pGro7.In Fig. 8, CDPV1 coexpressed with the molecular chaperone pTf16 had higher protein expression of the target protein at 25°C than at 18°C and 30°C, so the conditions at 25°C were chosen for subsequent experiments. Screening for IPTG concentrations was performed on proteins, as shown in Figure 9, where the target protein was induced to be more protein-expressed with 0.25mM IPTG than with 0.5mM IPTG at 25°C, so 0. 25mM IPTG was selected for induction and CDPV2-pTf16 and CDPV2-pGro7 were characterized under the same conditions, as shown in Figure 11, where CDPV2-pTf16 was more supernatantly expressed than CDPV2-pGro7, so CDPV2-pTf16 was selected.The subsequent protein purification was optimized from the induction conditions to determine that the target proteins obtained would be induced using CDPV1-pTf16 and CDPV2-pTf16 at 25 °C with 0.25mM of IPTG

**Figure 7.**
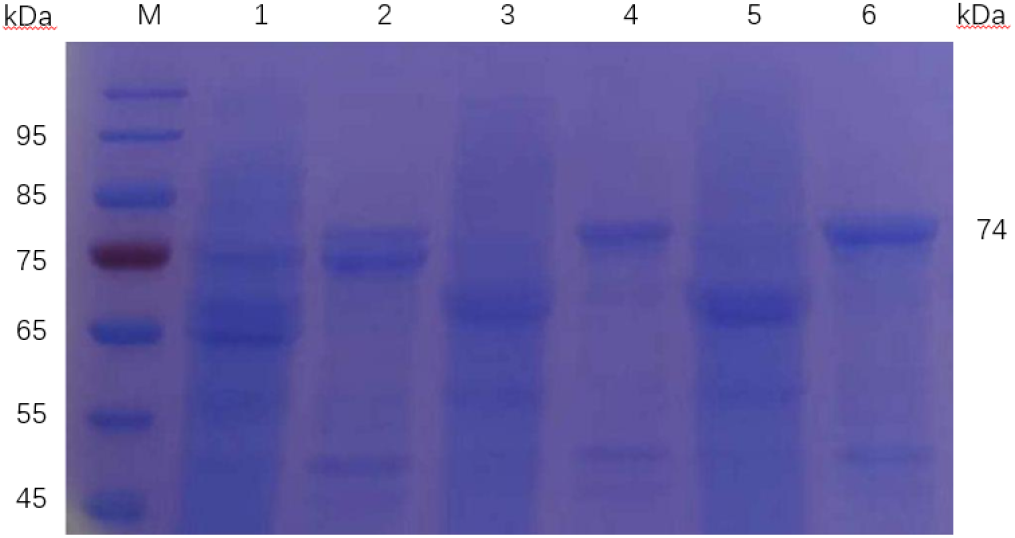
SDS-PAGE identification of CDPV1 cotransformed proteins with the molecular chaperone pGro7 at different temperatures. M:Maker; 1:30 °C CDPV1 coexpressed pGro7 supernatants; 2:30 °C CDPV1 coexpressed precipitates; 3:18 °C CDPV1 coexpressed p Gro7 supernatants; 4:18 °C CDPV1 coexpressed precipitates; 5:25 °C CDPV1 coexpressed p Gro7 supernatants; 6:25 °C CDPV1 coexpressed precipitates.

**Figure 8.**
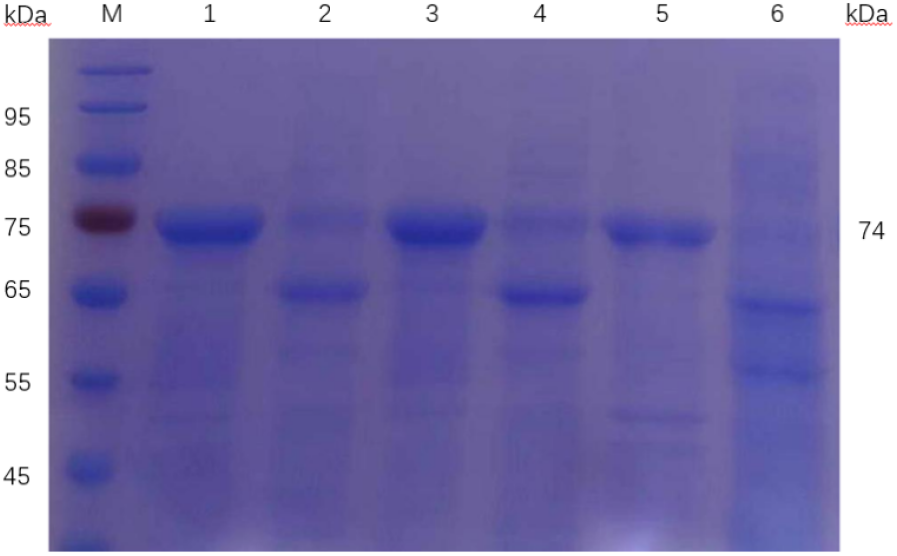
SDS-PAGE identification of CDPV1 cotransformed proteins with the molecular chaperone pTf16 at different temperatures. M:Maker;1:30 °C CDPV1 coexpressed pT f16 supernatants;2:30 °C CDPV1 coexpressed precipitates;3:18 °C CDPV1 coexpressed pTf16 supernatants;4:18 °C CDPV1 coexpressed precipitates;5:25 °C CDPV1 coexpressed pTf16 supernatants;6:25 °C CDPV1 coexpressed precipitates.

**Figure 9.**
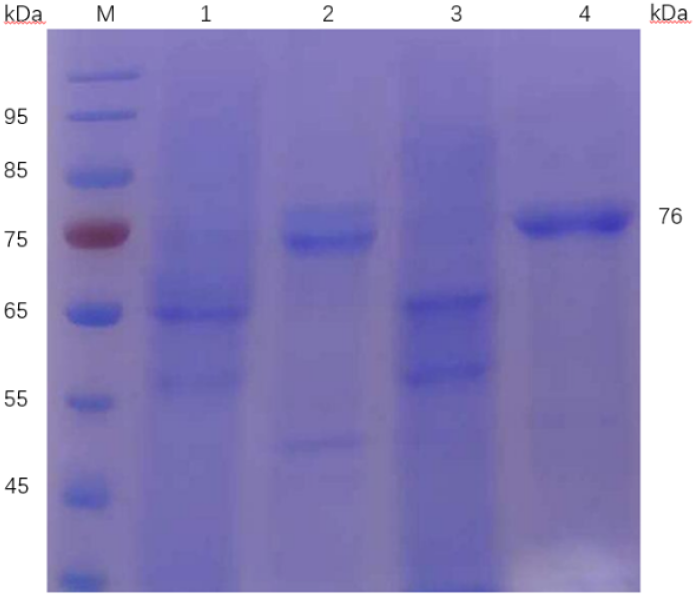
SDS-PAGE identification of CDPV1 cotransformed proteins with the molecular chaperone pTf16 at different IPTG concentrations. M.Maker;1:0.5 mM IPTG CDPV 1 co-expressed precipitate with pTf16;2:0.5 mM IPTG CDPV1 co-expressed supernatant with pTf16;3:0.25 mM IPTG CDPV1 co-expressed supernatant with pTf16,4:0.25 mM IPTG CDP V1 co-expressed supernatant with pTf16

**Figure 10.**
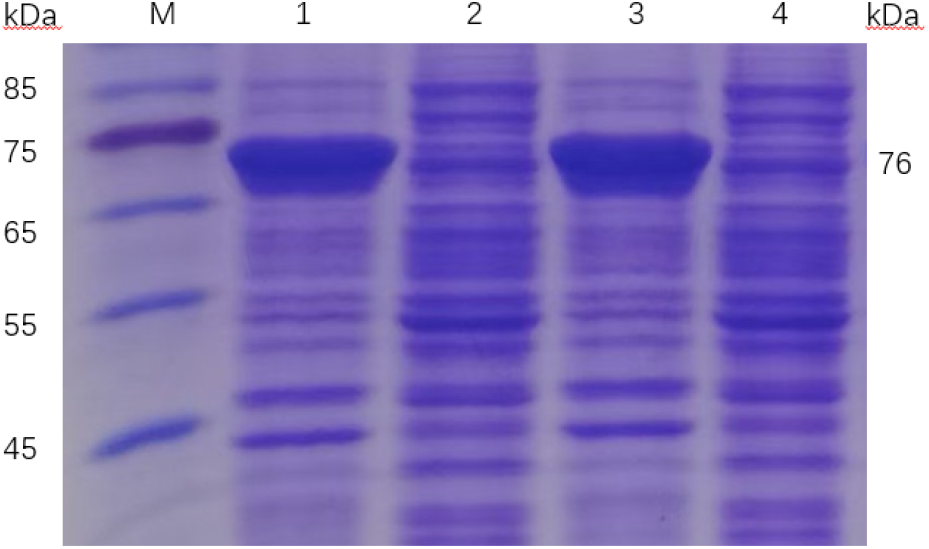
SDS-PAGE identification of CDPV2 and co-transformed proteins of the molecular chaperone pTf16, pGro7 M.Maker; 1: CDPV2 and pGro7 co-express supernatant s; 2: CDPV2 and pGro7 co-express pellets; 3: CDPV2 and pTf16 co-express supernatants; 4: CDPV2 and pTf16 co-express pellets;

**Figure 11.**
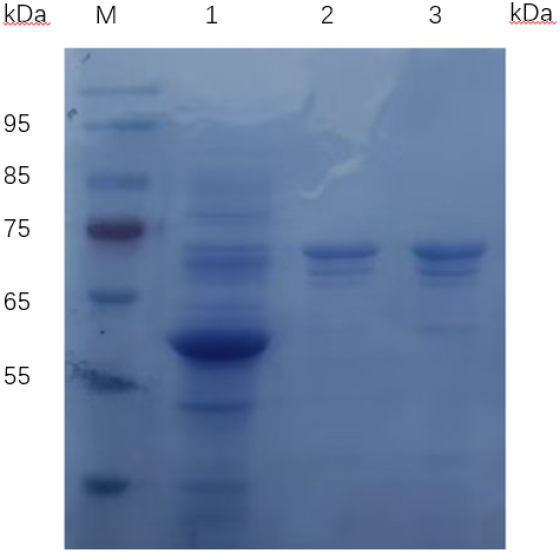
SDS-PAGE identification results of purified coexpressed proteins. M.Make r;1. Flow-through;2. Purified Protein C DPV1-pTf16;3. Purified Protein C DPV2-pTf16

**Figure 12.**
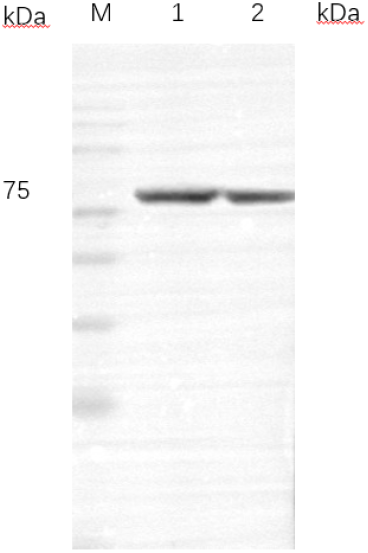
Western Blot identification results of purified coexpressed proteins. M.Maker;1. CDPV1-pTf16 ;2. CDPV2-pTf16

**Figure 13.**
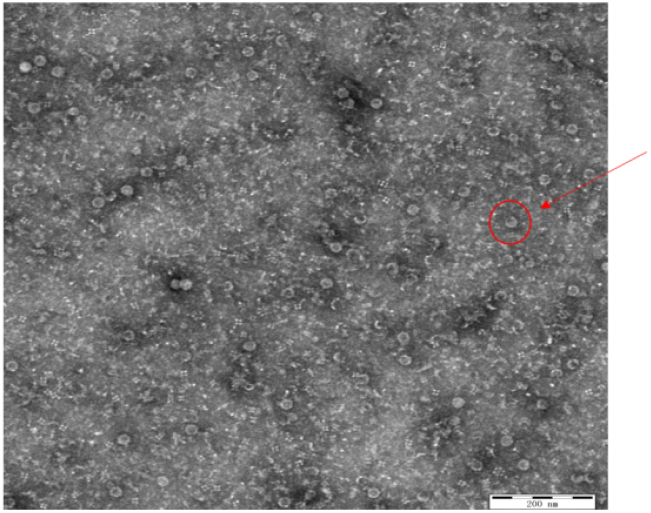
Recombinant protein transmission electron microscopy observation results. Scale bar: 200nm

#### Purification of recombinant proteins

The proteins obtained after fragmentation were purified by affinity nickel column purification, and purified after gradient elution. The purification effect was verified by SDS-PAGE. The results shown in Figure 11 showed that the proteins eluted with 150mM concentration of imidazol were the purest and free of obvious heteroproteins. Western Blot validation of the purified proteins showed that specific immunoblots were obvious at 74,76kDa.

#### Preparation and morphological observation of virus-like particles

The purified protein was put into a dialysis bag and allowed to assemble on its own in buffer and assembly solution. After overnight assembly, the resulting samples were centrifuge d and subjected to transmission electron microscopy. The TEM results are shown in Figure 1 3, which showed the formation of VLPs in spherical particles in the electron microscope, which were similar to natural CPV virions.DLS analysis further confirmed that the recombinant protein had a homogeneous particle size distribution, and as shown in Figure 14, the size of the particles was 23.5nm, close to the size of native CPV virions, indicating that the VLP successfully self-assembled to form a complete particles, indicating that the insertion of a suitable epitope in the VP2 protein did not affect the formation of the particles.

**Figure 14.**
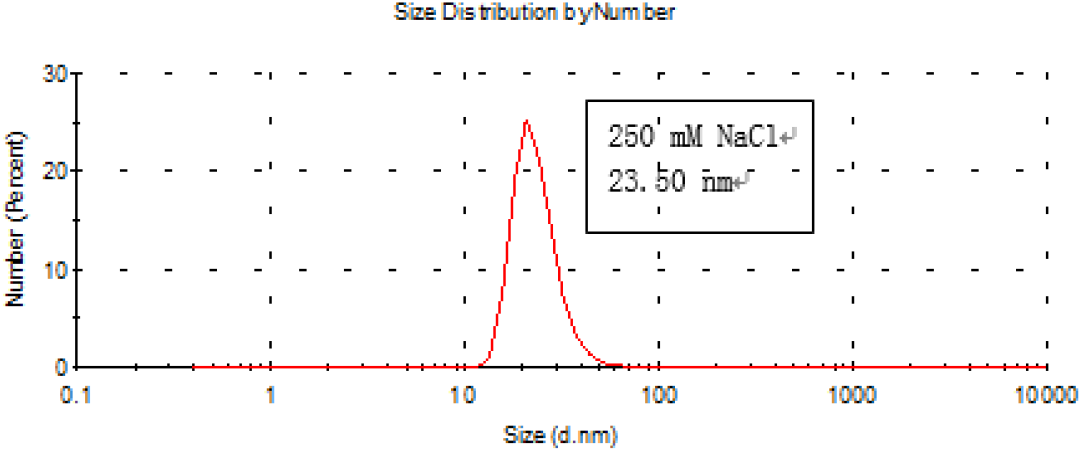
Dynamic light scattering results of recombinant proteins.

#### Detection of hemagglutination activity of virus-like particles

Given that natural CPV possesses porcine red blood cell, RBC agglutination activity (HA activity) at 4°C, the prepared samples were tested by hemagglutination (HA) test to assess their biological activity.The test results are shown in Figure 15, with a hemagglutination titer of 1:29 for both CDPV1 and CDPV2.This result suggests that the prepared virus-like particle s (VLPs) have similar porcine Red Blood Cell, RBC agglutination activity to natural CPV particles.

**Figure 15.**
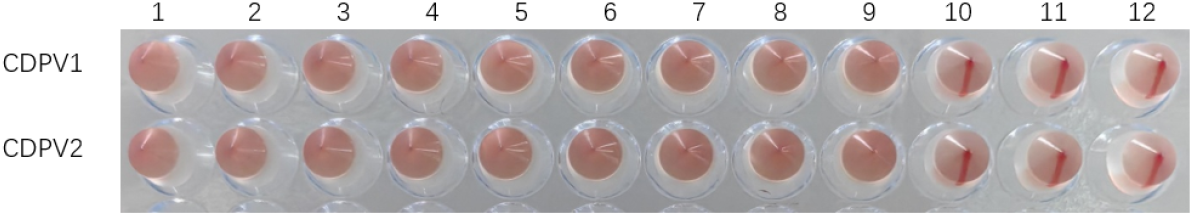
Results of the hemagglutination potency determination of CDPV1 and CDPV2.

#### Determination of the hemagglutination inhibition effect of CDPV1 and CDPV2 on mouse blood samples

Weekly blood samples were tested, and the results shown in Figure 16 showed that the peak hemagglutination inhibitory titer reached on day 7 after the third immunization in all groups of immunized mice; the peak antibodies were highest in the dual inactivated vaccine, followed by CDPV1+Al(OH)_3_, CDPV2 Al(OH)_3_, CDPV1, and CDPV2; CDPV1+Al(OH)_3_ and CDPV2+Al(OH)_3_ had increased hemagglutination inhibitory titers relative to CDPV1 and CD PV2 without AL(OH)_3_ addition.

**Figure 16.**
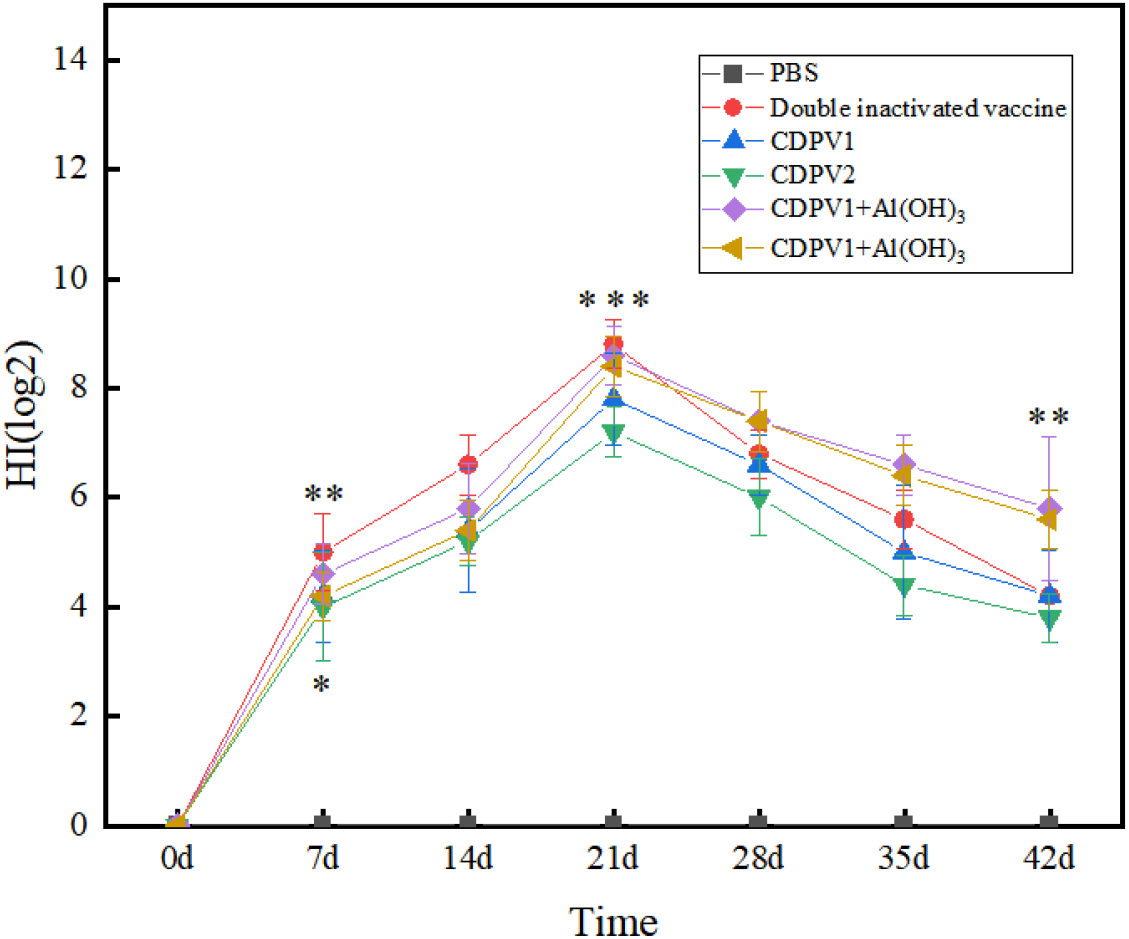
Trends in hemagglutination inhibition. \**P*<0.05, \*\**P*<0.01, \*\*\**P*<0.001 (t test).

#### Neutralizing effect

Neutralization test is one of the important indicators for evaluating the effectiveness of the vaccine. Serum from mice on the 7th day after the third immunization was collected and tested using microtiter neutralization test.As shown in Figure 17 and Figure 18, all other immune groups produced a certain amount of neutralizing antibodies except PBS. The highest levels of neutralizing antibodies were induced in the mice administered with the inactivated dual vaccine, which was approximately 1:2^8^. The levels of neutralizing antibodies induced in the mice administered with CDPV1+Al(OH)_3_ adjuvant and CDPV2+Al(OH)_3_ adjuvant were higher than those in the CDPV1 and CDPV2 groups.

**Figure 17.**
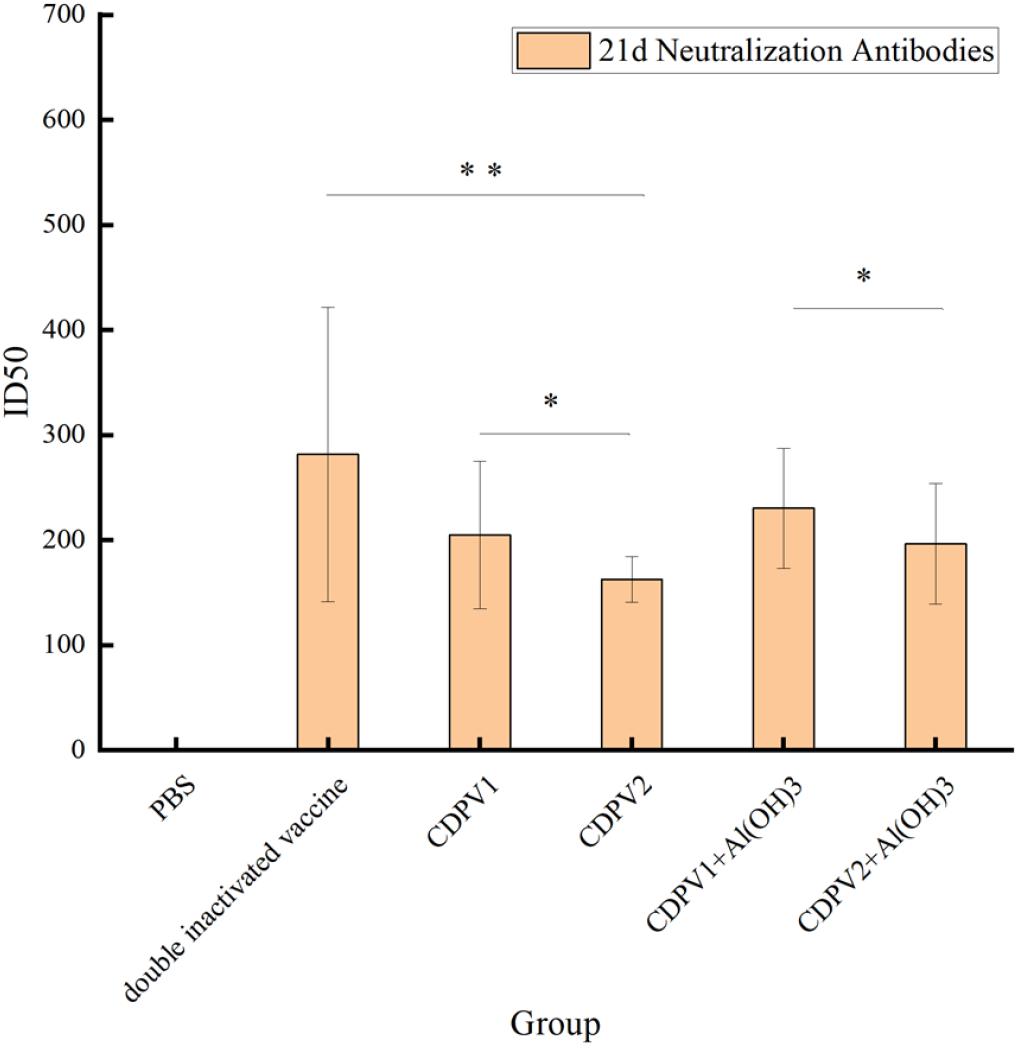
Detection of neutralizing Antibodies of CDV. \**P*<0.05, \*\**P*<0.01, \*\*\**P*<0. 001 (t test).

**Figure 18.**
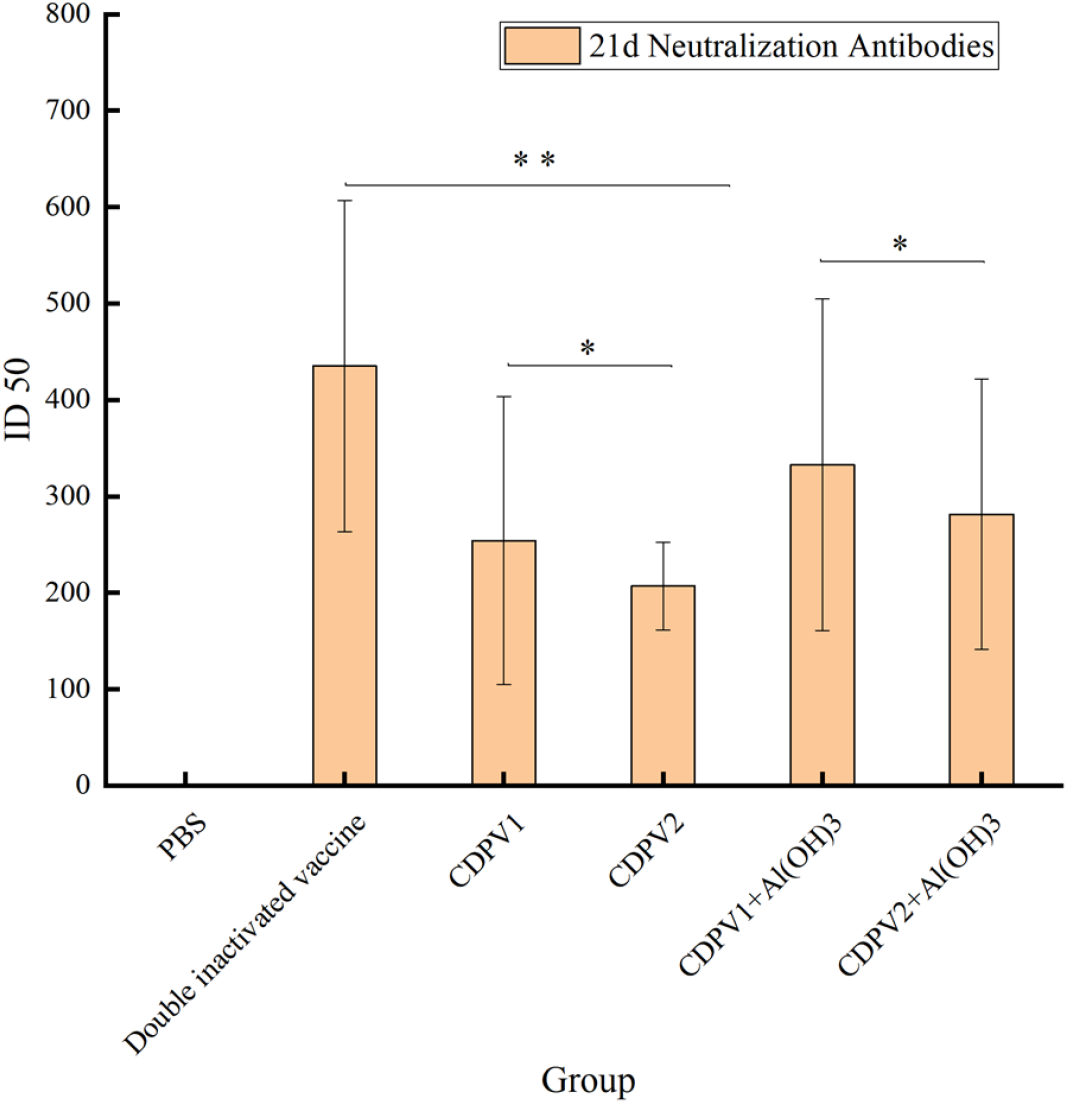
Detection of neutralizing Antibodies of CPV. \**P*<0.05, \*\**P*<0.01, \*\*\**P*<0. 001 (t test).

## Discussion

VLPs retain the antigen epitope and structural morphology of the natural viral particles and effectively present glycoproteins and other antigen components to the surface of the viral particles ^[10][13]^.As a particulate antigen, VLPs can mimic the in vivo infection process of wild-type viruses, effectively activate antigen-presenting cells (APCs), including Monocytes, macrophages, and DCs (DCs), and activate T cells by recognizing MHC class I molecules, leading to widespread appeal for the development of new subunit vaccines due to the high immunogenicity and safety of VLPs ^[14][16]^.

Studies have shown that CPV-VP2 can self-assemble in vitro to form a morphology and structure similar to that of a natural virus ^[17]^, Rueda ^[18]^, insertion of poliovirus C3 into four loops of CPV-VP2, found that epitope insertions in the four loops allowed for recapture of capsid in all mutants, where the N-terminal and Loop 2 regions of CPV-VP2 are nonessential regions in the VLP assembly process, and insertion of an exogenous antigen epitope between 218-233 amino acids in the Loop2 region of CPV-VP2 to construct chimeras did not affect t he in vitro formation of VLPs.

N, F, and H proteins in CDV are the major structural proteins of the virus ^[19]^.Zhao ^[20]^ et al. inserted epitopes on the F and N proteins of CDV in two combined ways into the N-terminal and Loop2 regions of CPV-VP2, obtained two recombinant plasmids, and enabled the m to successfully self-assemble and express.Studies on the prevalence of neutralizing antibodies against CDVs by TuBa^[21]^, et al., confirmed that the domain of H protein plays a key role in the cross-species transmission of CDVs, so we chose to insert antigen epitopes from the F, N, and H proteins of CDVs into CPV-VP2 to obtain recombinant plasmids.

Therefore, in this trial, a conservative prediction of epitopes in the literature was made t o demonstrate the feasibility of epitopes in this trial, and epitopes were evaluated by the web site to screen out those that could be used.Wang ^[22]^ et al. screened the H protein region of CDV for epitopes through the IEDB Analysis Resource (immuneepitope.org) website, and obtained available epitopes.Thus, by screening H proteins, we obtained the most antigenicity epitopes and utilized the structural plasticity of CPV-VP2 proteins to insert and fuse major B cel l epitopes, CTL epitopes, and T cell epitopes of CDVs, which formed chimeric CPV VLPs at the N terminus and Loop 2 region of CPV-VP2 proteins and displayed these epitopes on the surface of the CPV-VP2 VLPs.Finally, the separation of the synthetic recombinant plasmid and its three-dimensional structure were predicted. The solubility predictions of the recombinant plasmid reached 0.233 and 0.251, respectively, and the predicted results were insoluble expression, so the addition of molecular chaperones to the protein was considered to increase its solubility. The three-dimensional structure prediction scores were 0.77 and 0.78, respectively, indicating that the recombinant protein is stable in spatial structure.

The results of this electron microscopy test showed that VLP was successfully self-assembled at 250 mM/L NaCl with a pH of 8.0, and a VLP particle of 23.5 nm in size was obtained, which was similar to that of a natural canine parvovirus ^[23]^.The hemagglutination titer obtained for hemagglutination detection of the particles was 1:2^9^, consistent with the results of previous studies by this group ^[24]^, demonstrating that self-constructed recombinant plasmids have hemagglutination activity similar to that of natural CPV.In animal experiments, mice we re immunized with BALB/c mice by intramuscular injection of prepared CDPV1 and CDPV2 with Al(OH)_3_ adjuvants, and Serum was collected to detect the hemagglutination inhibition effect on mice blood samples. The results showed that both CDPV1 and CDPV2 had hemagglutination inhibition effects on mice blood samples.Neutralization results also showed that both CDPV1 and CDPV2 mice were able to induce a certain level of neutralizing antibodies, wit h the Al(OH)_3_ adjuvant group being higher than the no adjuvant group at 1:2^8^, and the CDP V1 group being higher than the CDPV2 group.It is illustrated that CDPV1 can induce stronger levels of neutralizing

Antibodies. In this study, recombinant Fusion Protein of CPV-VP2 with CDV antigen epitope was prepared to construct VLP vaccine candidates against CPV and CDV.In E.coli, the target protein was coexpressed with two chaperone proteins, and both chaperones were found to promote the soluble expression of the target protein. VP2 proteins were purified by ammonium sulfate precipitation and hydrophobic chromatography and successfully self-assembled. Th e virus-like granule vaccines CDPV1 and CDPV2 constructed showed good immunogenicity, which laid the foundation for the research of CDV and CPV prophylactic vaccines.

## Conclusion

This study successfully predicted and screened the antigenic epitopes of CDV proteins and successfully inserted them into CPV-VP2 proteins, constructing two virus-like granular vac cines, CDPV1 and CDPV2, which can successfully self-assemble in vitro and have good immunogenicity.The foundations were laid for the study of the CDV versus CPV dihybrid virus-like granular vaccines.

## Acknowledgments

We would like to extend our sincere gratitude to Changchun Sinor Co., Ltd. Additionally, we appreciate Xintao Liu from changchun BCHT Biotechnology Co.,Ltd.., for his help in serum sample processing.

## Author contributions statement

Data curation, Y.Z.and C.W. ; Formal analysis, Y.Z., G.Z. and Y.X.; Funding acquisition, C.W.; Investigation, Y.Z.; Methodology, Y.Z.; Project administration, C.W.; Resources, C. W.; Software, X.L.and Y.Z.; Supervision, C.W. and Y.Y.; Validation, Y.Y. and C.W.; Visualization,Y.Z., Y.X. and G.Z.; Writing—original draft,Y.Z., G.Z. and Y.X; Writing—review and editing, C.W. and Y.Y. All authors have read and agreed to the published version of the manuscript.

## Additional information

### Institutional review board statement

The animal study protocol was approved by the Animal Ethics Committee of Changchun Long Sheng Experimental Animal Technology Co., Ltd. (Approval Code: CCLSLL-2025082 817; Date: 28 August 2025), and the experimental operations were subject to the “Guidelines for the Welfare and Ethics of Experimental Animals in China”.

### Data availability statement

The data presented in this study are contained within the article.

